# Cardiac Outflow tract septation defects in a DiGeorge syndrome model respond to Minoxidil treatment

**DOI:** 10.1101/2024.03.18.585569

**Authors:** Ilaria Aurigemma, Rosa Ferrentino, Varsha Poondi Krishnan, Olga Lanzetta, Claudia Angelini, Elizabeth Illingworth, Antonio Baldini

## Abstract

**Background:** The T-BOX transcription factor TBX1 is essential for the development of the pharyngeal apparatus and it is haploinsufficient in DiGeorge syndrome (DGS), a developmental anomaly associated with congenital heart disease and other abnormalities. The murine model recapitulates the heart phenotype and showed collagen accumulation.

**Methods:** We first used a cellular model to study gene expression during cardiogenic differentiation of WT and *Tbx1*^-/-^ mouse embryonic stem cells. Then we used a mouse model of DGS to test whether interfering with collagen accumulation using an inhibitor of lysyl hydroxylase would modify the cardiac phenotype of the mutant.

**Results and conclusions:** In the cell differentiation model, loss of *Tbx1* was associated with up regulation of a subset of ECM-related genes, including several collagen genes. In the *in vivo* model, early prenatal treatment with Minoxidil, a lysyl hydroxylase inhibitor, ameliorated the cardiac outflow tract septation phenotype in *Tbx1* mutant fetuses, but it had no effect on septation in WT fetuses. We conclude that TBX1 suppresses a subset of ECM-related genes. The partial rescue of the septation phenotype through Minoxidil treatment suggests that inhibiting collagen cross-linking reduces the impact of the phenotypic consequences of *Tbx1* mutation.

## INTRODUCTION

Rescue experiments of disease gene phenotypes are important not only because they offer a potential avenue for treatment but also because they provide critical information about the importance of certain processes during normal development and the pathogenesis of the disease. However, phenotypic rescue may occur for different reasons including, but not limited to compensation for perturbed molecular pathways. *Tbx1* is a T-box transcription factor that is required for the development of the pharyngeal apparatus (Baldini et al., 2017) and in humans its haploinsufficiency causes a phenotypic spectrum known as DiGeorge syndrome, which is part of a broader clinical entity defined by a segmental aneuploidy known as 22q11.2 deletion syndrome (Paylor et al., 2006; Yagi et al., 2003). In the mouse, the *Tbx1* mutant phenotype, and particularly the cardiac phenotype, can be partially rescued using different drug-based and genetic approaches (Caprio and Baldini, 2014; Fulcoli et al., 2016; Lania et al., 2016; Racedo et al., 2017). The diversity of the approaches that have been at least partially successful suggests that there are a number of processes involved, correction of each of which individually is insufficient to resolve the developmental anomaly. In this study, we followed an alternative avenue and leveraged the findings of excess collagen that has been found in *Tbx1*^*-/-*^ embryos (Alfano et al., 2019).

We first used transcriptional analysis of a cell differentiation model and observed the upregulation of multiple collagen-encoding genes. Next, we used a drug-based approach *in vivo* to counterbalance the excessive collagen deposition observed in *Tbx1* mutant embryos (Alfano et al., 2019). A significant improvement in cardiac septation was observed, highlighting the importance of the ECM in this key developmental process and thereby justifying a more systematic dissection of the molecular pathways involved. Minoxidil, an inhibitor of lysyl hydroxylase, reduces collagen cross-linking (Murad and Pinnell, 1987) and it has been used successfully to ameliorate the thymic phenotype in an *in vitro* model of thymic hypoplasia caused by *Tbx1* mutation (Bhalla et al., 2022). Therefore, we tested *in vivo* treatment of *Tbx1* mutants with Minoxidil and found a significant improvement in cardiac septation.

## RESULTS AND DISCUSSION

### Loss of Tbx1 is associated with up regulation of a subset of ECM-related genes

We performed *in vitro* differentiation of mES cells that have been gene-edited to lack any functional *Tbx1* and the parental cell line (Cirino et al., 2020). We used the protocol published by Andersen et al. (2018) with minor modifications because it induces robust expression of the *Tbx1* gene at day 6 (d6) of differentiation (Andersen et al., 2018). Principal component analyses of the transcriptomes of the 4 biological replicates/genotype showed clear separation of WT and mutant samples (Fig. 1a). We generated a list of differentially expressed genes (DEGs) between *Tbx1*^-/-^ and *Tbx1*^+/+^ (parental cell line), calculated using NOISeq (Tarazona et al., 2015) and selecting a posterior probability of at least 0.95 and a fold change of at least 1.5 (Fig. 1b). We found 2033 DEGs, of which 1104 were down regulated in mutant cells and 929 were up regulated (genes are listed in the Suppl. Tab. 1). Gene Ontology analyses using the Cellular Component subset (GO:CC, highlighted in yellow in Suppl. Tab. 2) indicated that within the down regulated gene set, there was enrichment of terms related to muscle and heart development (Fig. 1c); in contrast, using the up regulated gene set we observed enrichment of terms related ECM (Fig. 1d, top 5; full results shown in Suppl. Tab. 2), genes related to these terms included several encoding collagens (Suppl. Tab. 2). Therefore, the results obtained in the cellular model were consistent with the finding of increased collagen expression in *Tbx1* mutant embryos (Alfano et al., 2019) and they extended those observed by Alfano et al. to additional ECM-related genes, suggesting a broader perturbation of the ECM compartment than previously suspected. We selected a group of 21 up regulated ECM-related genes (suggested by the gene ontology search and listed in Suppl. Tab. 3) and mapped the ENCODE cis candidate regulatory elements (cCREs) (Moore et al., 2020) to these genes. In total, we selected 318 regions that are listed in Suppl. Tab. 3. We next carried out a search for transcription factor (TF) binding motifs using HOMER (Heinz et al., 2010) and the mouse genome as the background. We found a number of motifs (the top 20 are shown in Suppl. Fig. 1), particularly motifs of the AP-1 complex of transcription factors, NF1, and TCF/bHLH factors. However, we did not find T-BOX binding motifs, suggesting that this subset of ECM-related genes may not be direct target of TBX1.

**Figure 1.**
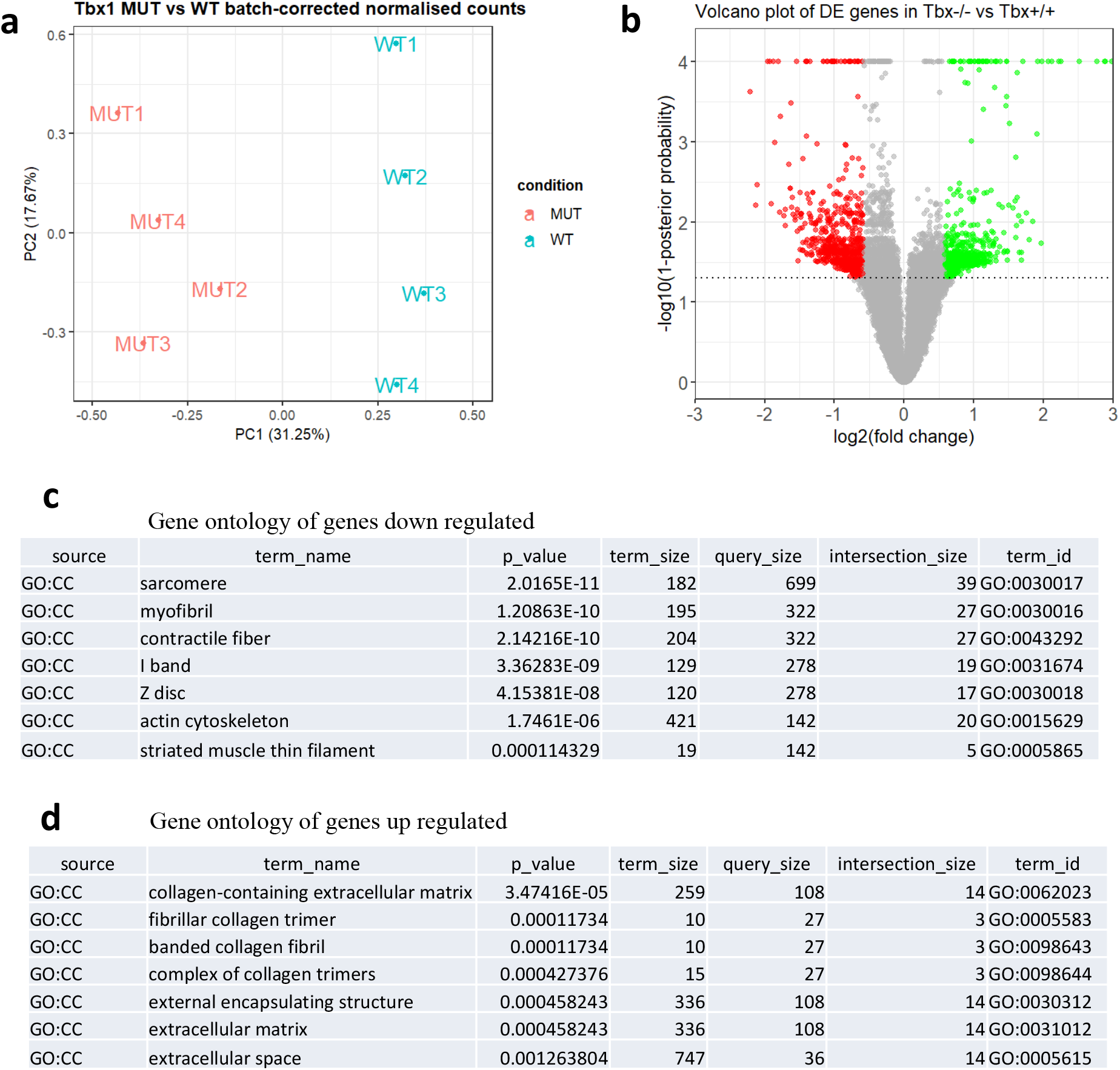
Gene expression analysis of cells differentiated using a cardiogenic protocol. a) Principal component analysis of QuantSeq data from 4 WT and 4 Tbx1-/-biological replicates (independent differentiation experiments). Samples are clearly separated by genotype. b) Volcano plot of gene expression values showing populations of differentially expressed genes. The individual values and gene names are listed in Suppl. Tab. 1 c-d) Top Gene Ontology / Cellular Component (GO:CC) terms identified for down regulated genes (c) and up regulated genes (d). Note the enrichment in ECM terms in the latter group. Full results with gene names are reported in Suppl. Tab. 2.

### Minoxidil treatment reduces the penetrance of outflow septation defects in Tbx1 mutants

An excess of ECM gene expression might impact cell dynamics/regionalization and the diffusion of signalling molecules, the developmental consequences of which are difficult to predict; cell regionalization abnormalities have been reported in *Tbx1* mutants (Lania et al., 2022). We asked whether counter-balancing the excess of ECM, particularly collagens, would modify the cardiac phenotype of *Tbx1* mutants. Addressing this question through genetic experiments would be very complex, due to the relatively large number of genes involved. Therefore, we opted for a pharmacological approach using a well-tolerated drug, Minoxidil, that inhibits lysyl hydroxylase. This destabilizes collagen by affecting cross-linking and reduces the accumulation of insoluble collagen (Rosell-García et al., 2019); this partially compensates for an excess of collagen. To this end, we crossed female mice carrying the hypomorphic *Tbx1* allele *Tbx1*^*neo2*/+^ (Zhang and Baldini, 2008) with *Tbx1*^*lacZ*/+^ male mice (the *lacZ* allele is a null allele (Lindsay et al., 2001)). We treated pregnant females with i.p. injections at E7.5, E8.5, and E9.5 with Minoxidil at 110µg/injection, or with vehicle (PBS+5% EtOH). Fetuses were harvested at E16.5 or E17.5, the hearts observed blind to genotype, and then processed for histology. Injection times were selected so as to target early SHF development when *Tbx1* is critical for OFT morphogenesis (Xu et al., 2005). We used the hypomorphic allele because *Tbx1*^*neo2*/lacZ^ fetuses have severe heart abnormalities but a less severe defective development of the pharyngeal apparatus (Zhang and Baldini, 2008). We harvested a total of 11 *Tbx1*^*neo2*/lacZ^ fetuses injected with vehicle (control) and found that 10 out of 11 had conal septation defects, resulting in a single, unseptated outflow valve (Tab. 1, Fig. 2a-d). Truncal septation was partial in 4 of these embryos as they showed incomplete separation of the aorta and pulmonary trunk (Tab. 1, Fig. 2c-c’). In only one case did we found double outlet right ventricle (DORV), which was associated with the complete separation of the aorta and pulmonary trunk, including the septation of the outflow valves (Fig. 2d-d’’). For purposes of comparison, the sections of a WT heart are shown in Fig. 2a-a’’). We harvested 13 *Tbx1*^*neo2*/lacZ^ fetuses injected with Minoxidil, of which 5 had septation defects of the outflow valve (Tab. 1, examples in Fig. 3a-e), while 8 had normally septated outflow valves. Five of the latter had DORV, while 3 had normal OFT morphology but a mild VSD (example in Fig. 3b). Septation defects of the OFT in *Tbx1*^*neo2*/lacz^ fetuses were significantly less prevalent after Minoxidil treatment (P=0.008) (Fig. 3g). We also examined 4 WT fetuses that were treated with Minoxidil and found no morphological anomalies (examples in Fig. 4a-c). Thus, Minoxidil treatment as described here does not interfere with OFT development in the WT.

**Figure 2.**
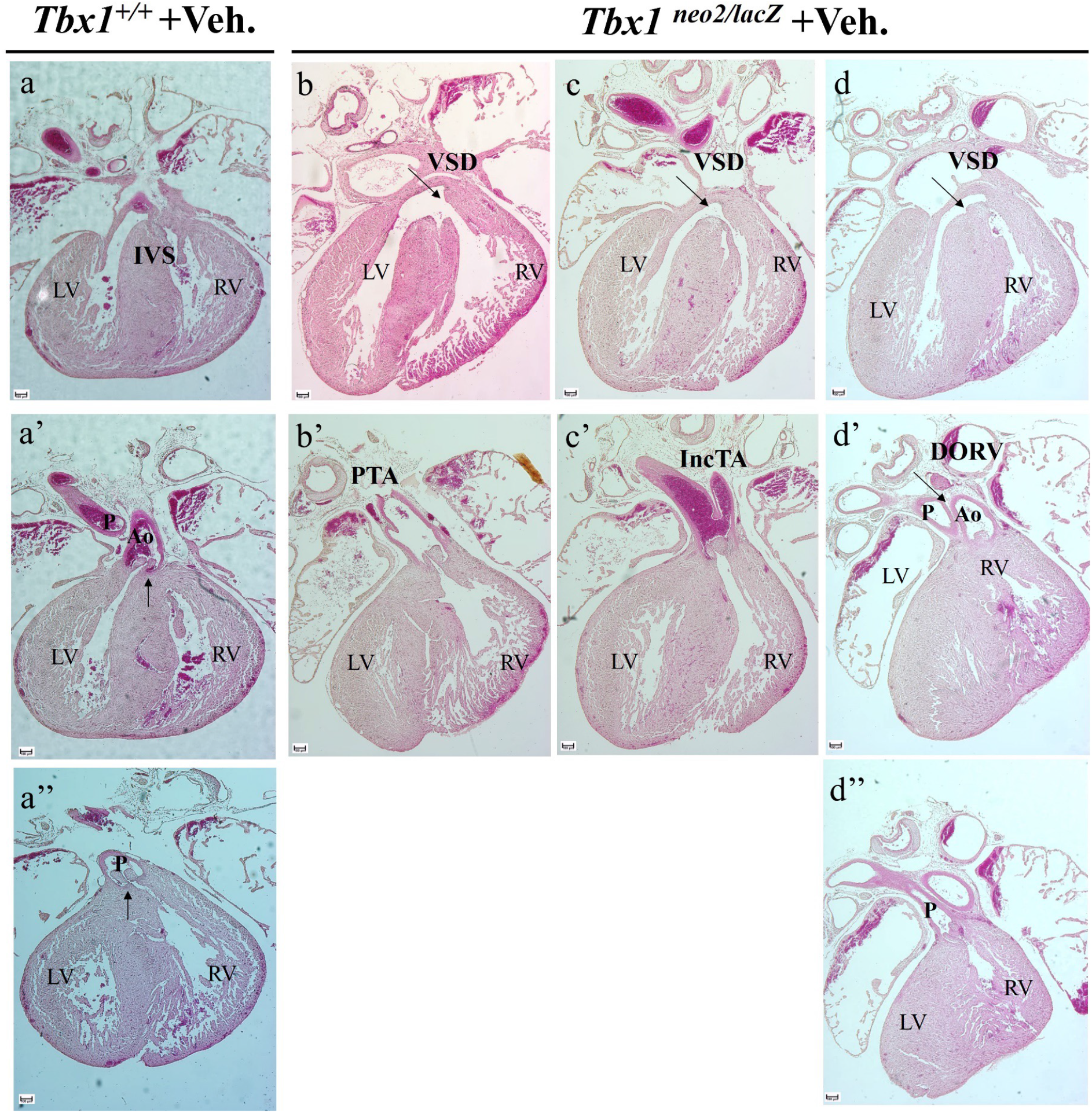
Cardiac outflow tract phenotype in vehicle-treated *Tbx1*^*neo2*/lacZ^ fetuses (E16.5-E17.5). a-a’’) H&E stained histological sections of a WT heart as reference. b-d) three examples of vehicle-treated *Tbx1*^*neo2*/lacZ^ fetuses. Sections indicated with the same letter are from the same heart. IVS: Interventricular septum; LV: Left ventricle; RV: Right ventricle; VSD: Ventricular septal defect; PTA: Patent truncus arteriosus; IncTA: Incomplete septation of the arterial trunk; Ao: Aorta; P: Pulmonary trunk; DORV: Double outlet right ventricle. Scale bare: 100µm.

**Figure 3.**
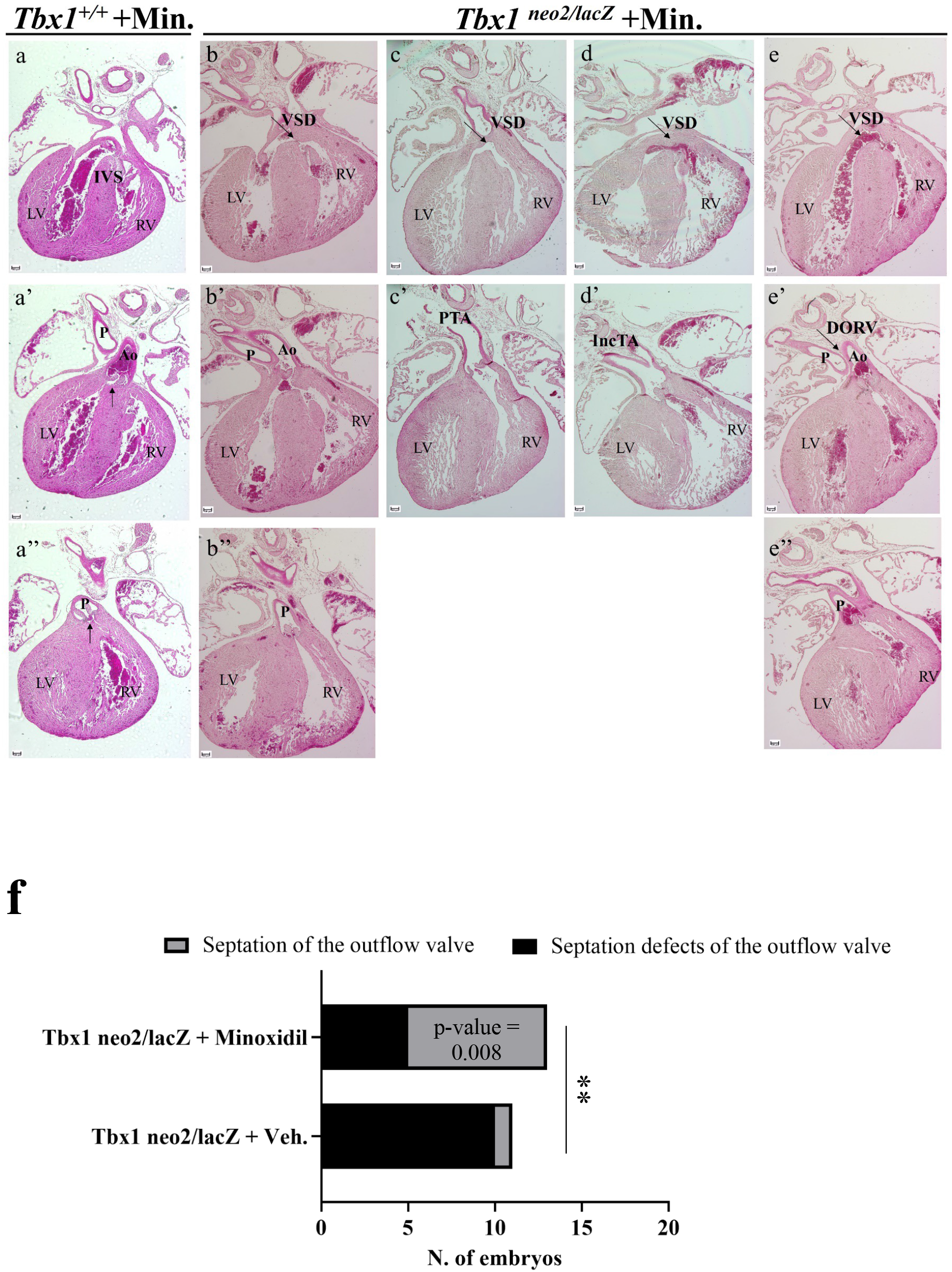
Cardiac outflow tract phenotype in Minoxidil-treated *Tbx1*^*neo2*/lacZ^ fetuses (E16.5-E17.5). a-a’’) H&E stained histological sections of a WT heart as reference. b-e) Examples of hearts from Minoxidil-treated *Tbx1*^*neo2*/lacZ^ fetuses. f) the incidence of septation defects (one OFT valve vs. two OFT valves) in Minoxidil-treated individuals is significantly lower than in vehicle-treated individuals (Chi-square test). IVS: Interventricular septum; LV: Left ventricle; RV: Right ventricle; VSD: Ventricular septal defect; PTA: Patent truncus arteriosus; IncTA: Incomplete septation of the arterial trunk; Ao: Aorta; P: Pulmonary trunk; DORV: Double outlet right ventricle. Scale bare: 100µm.

**Figure 4.**
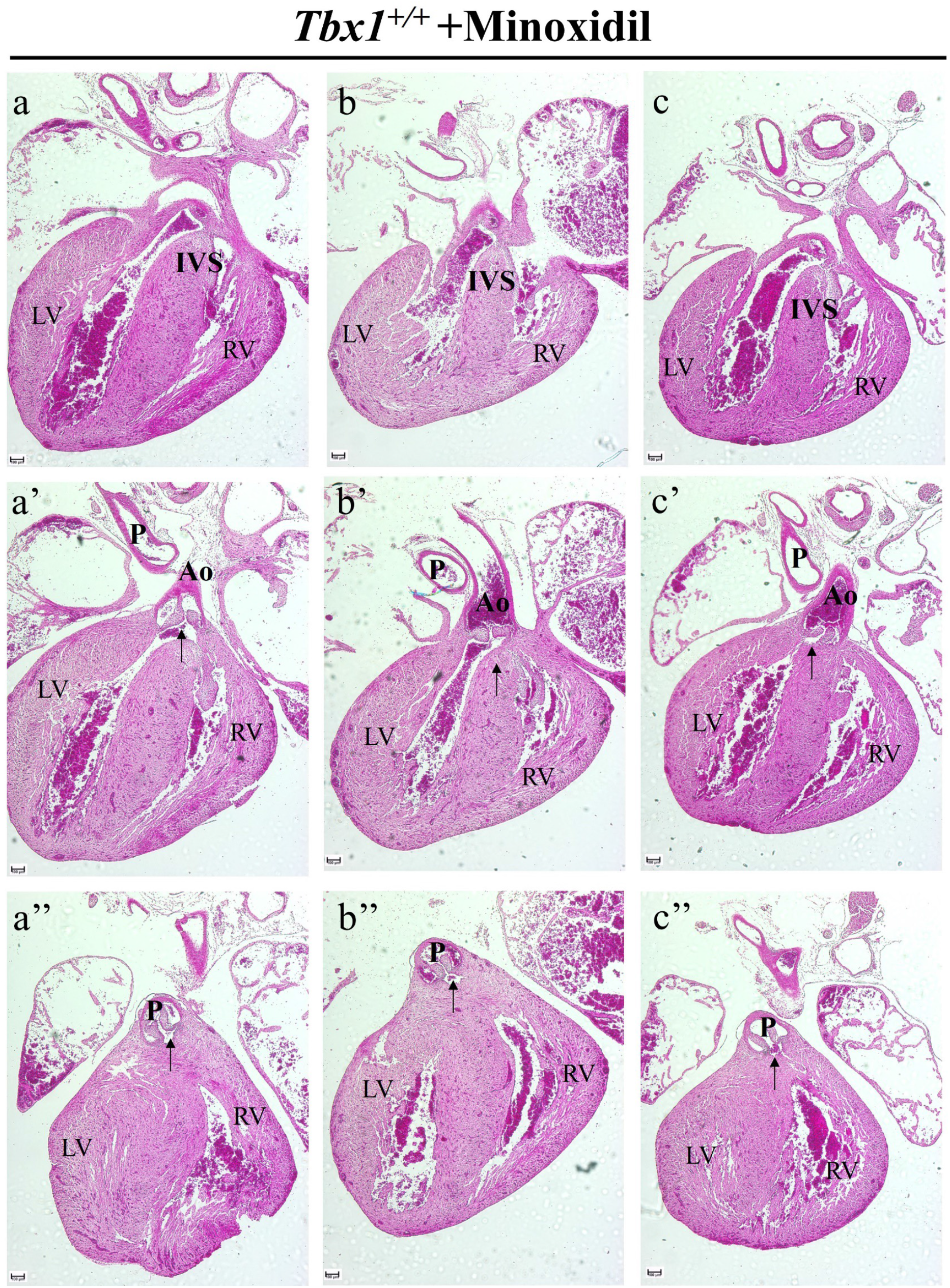
Minoxidil treatment does not affect normal development of the OFT in WT individuals. a-c) Hematoxylin & eosin-stained sections of hearts from fetuses WT fetuses treated with Minoxidil. There are no morphological defects. Sections indicated with the same letter are from the same heart. IVS: Interventricular septum; LV: Left ventricle; RV: Right ventricle; Ao: Aorta; P: Pulmonary trunk; Scale bare: 100µm.

These results indicate that Minoxidil treatment during SHF development is a significant modifier of the cardiac OFT septation phenotype in *Tbx1* mutants. Because the treatment was carried out well before the actual process of septation takes place, and because the drug, under the conditions tested, did not interfere with OFT formation, we propose that the drug interacts with or partially compensates for the alterations caused by *Tbx1* mutation. In addition to the cited role of the drug as a lysyl hydroxylase inhibitor, Minoxidil is an anti-hypertensive drug.

This effect is thought to be produced through relaxation of vascular smooth muscle by acting on ATP-sensitive potassium (KATP) channels of the sarcolemma (Wang, 2003). This particular mechanism is unlikely to be relevant in the context of our study because at the time of treatment, there is little or no smooth muscle differentiation in the cardiovascular system. Minoxidil is also used topically to treat androgenetic alopecia, but the mechanism by which it acts in this context is unclear (Suchonwanit et al., 2019). The finding that Minoxidil ameliorates the thymic phenotype in an *in vitro* model of thymic hypoplasia caused by *Tbx1* mutation (Bhalla et al., 2022) supports the hypothesis that it interferes with a pathological process triggered by reduced *Tbx1* gene dosage.

## Conclusions

We found that in mice, *Tbx1* suppresses a group of ECM-related genes including multiple collagen genes; *in vivo* treatment with a drug that reduces collagen cross-linking leads to significant improvement of cardiac outflow tract development in *Tbx1* hypomorphic mutants, suggesting that an excess of collagen may be part of the pathogenetic mechanism or modify significantly the phenotypic outcome of reduced gene dosage. Further work will be necessary to establish the molecular links through which *Tbx1* suppresses ECM-related genes.

## ACKNOWLEDGEMENTS

We thank Nicolai Van Oers and Pratibha Bhalla for suggestions about Minoxidil treatment and Marchesa Bilio for expert technical help. RNA-seq experiments were carried out at the Telethon Institute of Genetics and Medicine, Pozzuoli, Italy on a fee-for-service basis.

## COMPETING INTERESTS

‘No competing interests declared’.

## FUNDING

This work was funded in part by grants from Fondazione Telethon GMR22T1012 and the Italian Ministry of University and Research PRIN 2022XFE7M2 (to AB).

## DATA AVAILABILITY

All relevant data can be found within the article and its supplementary information.

RNA-seq data are available through the GEO database, accession number GSE261206.

## MATERIALS AND METHODS

### Cell culture and differentiation

The embryonic stem cell line 4D carries a homozygous targeted inactivating mutation of the *Tbx1* gene and it was obtained using CRISPR-Cas9 targeting (Cirino et al., 2020). This cell line was cultured along with the parental mouse ES cell line ES-E14TG2a (ATCC CRL-1821) without feeders, maintaining their undifferentiated state on gelatin-coated dishes. The culture medium consisted of GMEM (Sigma Cat# G5154) supplemented with 10^3^ U/ml ESGRO LIF (Millipore, Cat# ESG1107), 15% fetal bovine serum (ES Screened Fetal Bovine Serum, US Euroclone Cat# CHA30070L), 0.1 mM non-essential amino acids (Gibco, Cat# 11140-035), 0.1 mM 2-mercaptoethanol (Gibco, Cat# 31350-010), 0.1 mM L-glutamine (Gibco, Cat# 25030081), 0.1 mM Penicillin/Streptomycin (Gibco, Cat# 10378016), and 0.1 mM sodium pyruvate (Gibco, Cat# 11360-070). Cell passaging was performed every 2–3 days using 0.25% Trypsin-EDTA (1X) (Gibco, Cat# 25200056) as the dissociation buffer.

For differentiation, E14-Tg2a cells (parental line and *Tbx1*^-/-^ line 4D) were dissociated with Trypsin-EDTA and cultured at a density of 100,000 cells/ml in serum-free differentiation media comprising 75% Iscove’s modified Dulbecco’s media (Cellgro Cat# 15-016-CV) and 25% HAM F12 media (Cellgro #10-080-CV). The media were supplemented with N2 (GIBCO #17502048) and B27 (GIBCO #12587010) supplements, penicillin/streptomycin (GIBCO #10378016), 0.05% BSA (Invitrogen Cat#. P2489), L-glutamine (GIBCO #25030081), 5 mg/ml ascorbic acid (Sigma A4544), and 4.5 × 10-4M monothioglycerol (Sigma M-6145). After 48 hours, the embryoid bodies were collected and plated in serum-free differentiation media, supplemented with 1 ng/ml human Activin A (R&D Systems Cat#. 338-AC) and 1 ng/ml human BMP4 (R&D Systems Cat# 314-BP). The media were changed to serum-free differentiation media without additional growth factors after 2 days, and the embryoid bodies were maintained for 96 hours, completing the differentiation protocol.

### Gene expression analysis

For RNA-seq/QuantiSeq experiments, RNA was extracted from 4D and parental cell line at d6 of differentiation from 4 independent differentiation experiments (4 biological replicates) and processed for QuantiSeq 3’ mRNA sequencing. Libraries were prepared and sequenced at the Telethon Institute of Genetics and Medicine of Pozzuoli, Italy on a fee-for-service basis. We performed quality control and trimming, including polyA tails and adapter removal, with the bbduck.sh script from the BBTools suite (https://sourceforge.net/projects/bbmap/) with k=13 ktrim=r useshortkmers=t mink=5 qtrim=r trimq=10 minlength=20. Then, we aligned the sequences to the mouse genome (mm10) using STAR (Dobin et al., 2013), and obtained the gene expression as a raw count matrix using the featureCounts function from the Rsubread package. We filtered out genes with null or very low expression using the filtrered.data function from the NOISeq package (Tarazona et al., 2015) using the proportion test as the filtering method and retaining only the genes with counts per million (CPM) above 0.5, as expressed genes. We removed the batch using the ARSyNseq function in the same package with upper quartile normalization and visualized the quality of the samples in the principal component projection. To account for biological variability, we used the noiseqbio function, setting the cutoff for the posterior probability to 0.95, to identify the differentially expressed genes. Gene ontology of the differentially expressed genes was performed with the gprofiler online tool (https://biit.cs.ut.ee/gprofiler/gost) using the entire set of expressed genes as the background.

To collect putative regulatory elements of ECM genes, we downloaded all the cis candidate regulatory elements (cCREs) in mm10 from https://screen.encodeproject.org/ of the ENCODE project (Moore et al., 2020) and considered only distal enhancer-like sequences (dELS) and proximal enhancer-like sequences (pELS). Next, we used the getbioregion() function from ChIPseeker (v1.38.0) (Yu et al., 2015) to obtain the gene body coordinates using the TxDb.Mmusculus.UCSC.mm10.knownGene (v.3.10.0) database and we extended the gene body 50 Kb upstream of the transcription start site (TSS) and 50Kb downstream of the transcription end site (TES). We intersected the gene coordinates with the selected cCREs regions using the FindOverlaps function from GenomicRanges(v.1.54.1) (Lawrence et al., 2013) to retain only regions that overlap with the genes. We filtered the cCREs regions with annotated genes intersecting a list of 21 upregulated genes and we obtained a list of 318 regions (Supplementary Table 3). Finally, we utilized these regions to identify enriched motifs using HOMER (Hypergeometric Optimization of Motif EnRichment) (Heinz et al., 2010), setting as background the annotated cCREs regions that overlap with all the expressed genes (n= 16391, Supplementary Tab. 3) using the same procedure described above. We repeated the HOMER search using the default background (mouse genome).

### *In vivo* treatment with Minoxidil

Mouse lines *Tbx1*^*lacZ*^/+ (Lindsay et al., 2001) (also indicated here as *Tbx1*^+/-^) and *Tbx1*^*neo2*/+^ (Zhang and Baldini, 2008) have been maintained and used in a C57Bl/6N genetic background and genotyped according to the original description. Pregnant females were injected intraperitoneally with the vehicle (PBS with 5% EtOH) or Minoxidil (Sigma-Aldrich M4145). Each injection included 110µg of Minoxidil or vehicle in 200 µl, at E7.5, E8.5, and E9.5. Fetuses were harvested at E16.5 or E17.5, hearts were dissected, examined and embedded in paraffin for histology after fixation in 4% paraformaldehyde (PFA). Sections (10 µm) were stained with Hematoxylin and Eosin (Mayer′s Hematoxylin Solution MHS16, Eosin Y Solution, Alcoholic HT110116), using standard methods. Images were acquired using a Nikon Eclipse Ni with a 4x objective equipped with Nikon DS-RI1 Camera and software Nis Elements AR 4.20.00.

The rescue of septation defects of the OFT between treated and control *Tbx1*^*neo2*/lacZ^fetuses was tested using the Chi-square test.

All animal procedures were reviewed and approved by the Italian Ministry of Health, protocol 28/2022-2PR (to AB). Mouse lines are available through the European public repository EMMA/Infrafrontier.

## AUTHOR CONTRIBUTIONS

IA performed experiments, contributed to experimental design, edited the manuscript; RF performed experiments; OL performed data analysis; CA supervised data analyses; VPK performed data analysis, edited the manuscript; EI provided funding, edited the manuscript; AB provided funding, wrote the manuscript.

## FIGURE LEGENDS

**Table 1.**
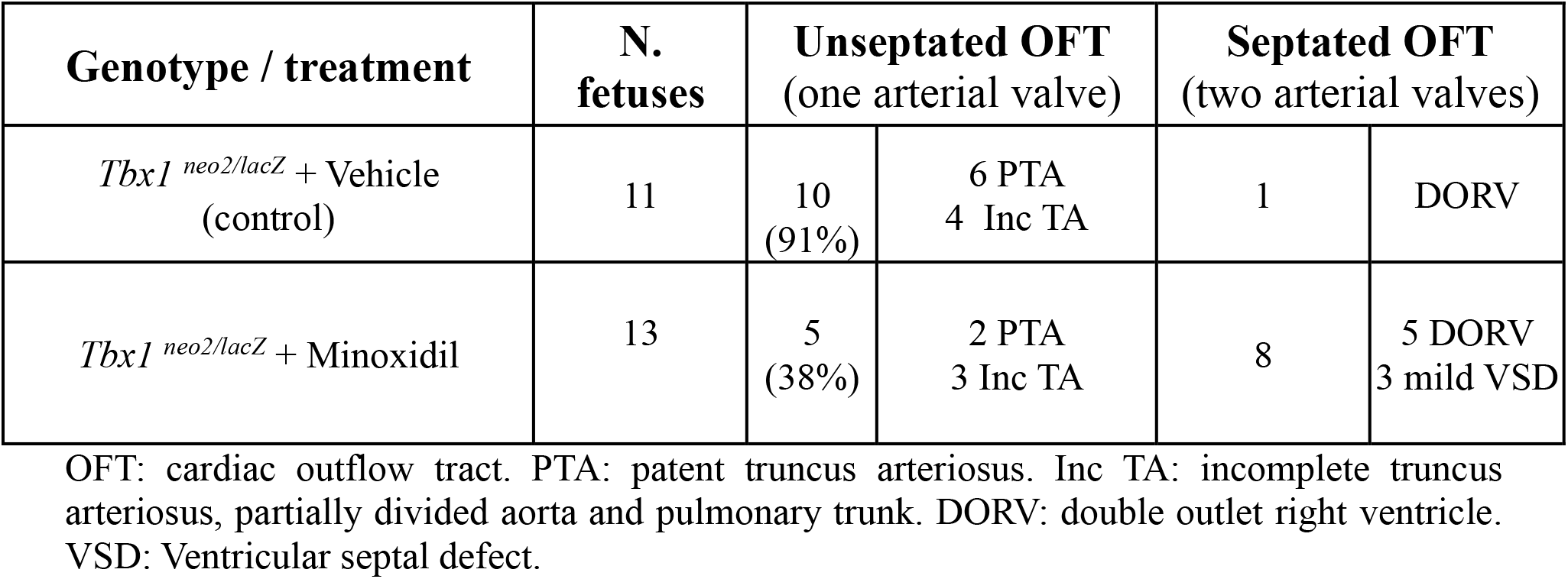
Summary of cardiac outflow septation status in control and Minoxidil-treated fetuses.

| **Genotype / treatment** | **N.**<br>**fetuses** | **Unseptated OFT**<br>(one arterial valve) |  | **Septated OFT**<br>(two arterial valves) |  |
| --- | --- | --- | --- | --- | --- |
| *Tbx1*<sup>neo2/lacZ</sup> + Vehicle<br>(control) | 11 | 10<br>(91%) | 6 PTA<br>4 Inc TA | 1 | DORV |
| *Tbx1*<sup>neo2/lacZ</sup> + Minoxidil | 13 | 5<br>(38%) | 2 PTA<br>3 Inc TA | 8 | 5 DORV<br>3 mild VSD |

OFT: cardiac outflow tract. PTA: patent truncus arteriosus. Inc TA: incomplete truncus arteriosus, partially divided aorta and pulmonary trunk. DORV: double outlet right ventricle. VSD: Ventricular septal defect.

## SUPPLEMENTARY MATERIAL

**Supplementary Figure 1**

HOMER output of 318 candidate regulatory elements mapped to the 21 up regulated genes related to extra cellular matrix, using the mouse genome as background.

**Supplementary Table 1**

Sheet 1: Differentially expressed genes (KO vs WT cells at d6 of differentiation)

Sheet 2: Genes UP regulated

Sheet 3: Genes DOWN regulated

**Supplementary Table 2**

Sheet 1: Gene Ontology search results using the down regulated genes.

Sheet 2: Gene Ontology search results using the up regulated genes.

**Supplementary Table 3**

Sheet 1: List of 21 genes upregulated and related to extra cellular matrix (ECM).

Sheet 2: List of ENCODE cis candidate regulatory elements mapped to the 21 ECM-related genes.

## Notes

### Competing Interest Statement

The authors have declared no competing interest.

