## Supplementary Figure 1 for "Cardiac Outflow tract septation defects in a DiGeorge syndrome model respond to Minoxidil treatment"

| Rank | Motif | Name | P-value | log P-pvalue | q-value (Benjamini) | # Target Sequences with Motif | % of Targets Sequences with Motif | # Background Sequences with Motif | % of Background Sequences with Motif | Motif File | SVG |
|---|---|---|---|---|---|---|---|---|---|---|---|
| 1 | 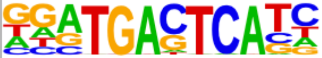 | Fra2(bZIP)/Striatum-Fra2-ChIP-Seq(GSE43429)/Homer | 1e-6 | -1.421e+01 | 0.0003 | 40.0 | 12.58% | 2646.9 | 5.38% | motif file (matrix) | svg |
| 2 | 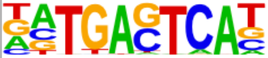 | BATF(bZIP)/Th17-BATF-ChIP-Seq(GSE39756)/Homer | 1e-5 | -1.364e+01 | 0.0003 | 50.0 | 15.72% | 3779.0 | 7.68% | motif file (matrix) | svg |
| 3 | 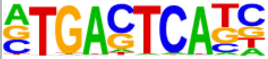 | AP-1(bZIP)/ThioMac-PU.1-ChIP-Seq(GSE21512)/Homer | 1e-5 | -1.305e+01 | 0.0003 | 54.0 | 16.98% | 4309.4 | 8.76% | motif file (matrix) | svg |
| 4 | 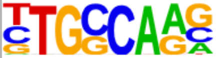 | NF1-halfsite(CTF)/LNCaP-NF1-ChIP-Seq(Unpublished)/Homer | 1e-5 | -1.234e+01 | 0.0005 | 117.0 | 36.79% | 12483.8 | 25.37% | motif file (matrix) | svg |
| 5 | 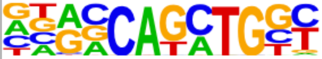 | Atoh1(bHLH)/Cerebellum-Atoh1-ChIP-Seq(GSE22111)/Homer | 1e-5 | -1.220e+01 | 0.0005 | 83.0 | 26.10% | 7984.4 | 16.22% | motif file (matrix) | svg |
| 6 | 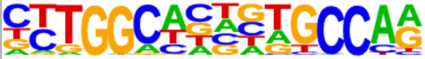 | NF1(CTF)/LNCAP-NF1-ChIP-Seq(Unpublished)/Homer | 1e-5 | -1.197e+01 | 0.0005 | 36.0 | 11.32% | 2482.7 | 5.04% | motif file (matrix) | svg |
| 7 | 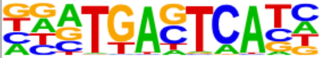 | Fra1(bZIP)/BT549-Fra1-ChIP-Seq(GSE46166)/Homer | 1e-5 | -1.174e+01 | 0.0005 | 41.0 | 12.89% | 3041.2 | 6.18% | motif file (matrix) | svg |
| 8 | 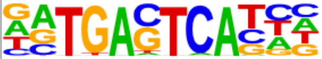 | Atf3(bZIP)/GBM-ATF3-ChIP-Seq(GSE33912)/Homer | 1e-4 | -1.148e+01 | 0.0006 | 47.0 | 14.78% | 3742.3 | 7.60% | motif file (matrix) | svg |
| 9 | 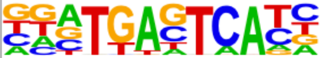 | Fos(bZIP)/TSC-Fos-ChIP-Seq(GSE110950)/Homer | 1e-4 | -1.126e+01 | 0.0006 | 42.0 | 13.21% | 3215.0 | 6.53% | motif file (matrix) | svg |
| 10 | 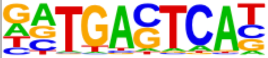 | JunB(bZIP)/DendriticCells-Junb-ChIP-Seq(GSE36099)/Homer | 1e-4 | -1.126e+01 | 0.0006 | 41.0 | 12.89% | 3104.8 | 6.31% | motif file (matrix) | svg |
| 11 | 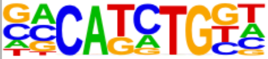 | TCF4(bHLH)/SHSY5Y-TCF4-ChIP-Seq(GSE96915)/Homer | 1e-4 | -1.116e+01 | 0.0006 | 106.0 | 33.33% | 11268.3 | 22.90% | motif file (matrix) | svg |
| 12 | 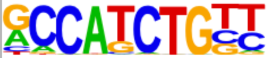 | NeuroD1(bHLH)/Islet-NeuroD1-ChIP-Seq(GSE30298)/Homer | 1e-4 | -1.076e+01 | 0.0008 | 64.0 | 20.13% | 5881.8 | 11.95% | motif file (matrix) | svg |
| 13 | 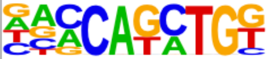 | BHLHA15(bHLH)/NIH3T3-BHLHB8.HA-ChIP-Seq(GSE119782)/Homer | 1e-4 | -1.019e+01 | 0.0013 | 97.0 | 30.50% | 10293.6 | 20.92% | motif file (matrix) | svg |
| 14 | 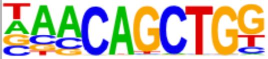 | Tcf21(bHLH)/ArterySmoothMuscle-Tcf21-ChIP-Seq(GSE61369)/Homer | 1e-4 | -9.414e+00 | 0.0026 | 70.0 | 22.01% | 6916.2 | 14.05% | motif file (matrix) | svg |
| 15 | 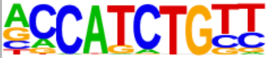 | NeuroG2(bHLH)/Fibroblast-NeuroG2-ChIP-Seq(GSE75910)/Homer | 1e-4 | -9.246e+00 | 0.0028 | 102.0 | 32.08% | 11228.3 | 22.82% | motif file (matrix) | svg |
| 16 | 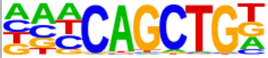 | Ap4(bHLH)/AML-Tfap4-ChIP-Seq(GSE45738)/Homer | 1e-3 | -8.186e+00 | 0.0077 | 84.0 | 26.42% | 9067.4 | 18.42% | motif file (matrix) | svg |
| 17 | 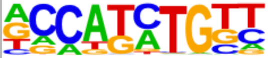 | Olig2(bHLH)/Neuron-Olig2-ChIP-Seq(GSE30882)/Homer | 1e-3 | -7.966e+00 | 0.0090 | 118.0 | 37.11% | 13868.6 | 28.18% | motif file (matrix) | svg |
| 18 | 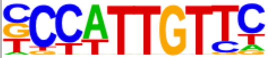 | Sox2(HMG)/mES-Sox2-ChIP-Seq(GSE11431)/Homer | 1e-3 | -7.418e+00 | 0.0147 | 55.0 | 17.30% | 5453.7 | 11.08% | motif file (matrix) | svg |
| 19 | 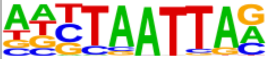 | Lhx3(Homeobox)/Neuron-Lhx3-ChIP-Seq(GSE31456)/Homer | 1e-3 | -7.322e+00 | 0.0153 | 92.0 | 28.93% | 10414.3 | 21.16% | motif file (matrix) | svg |
| 20 | 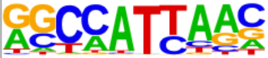 | Nanog(Homeobox)/mES-Nanog-ChIP-Seq(GSE11724)/Homer | 1e-3 | -7.259e+00 | 0.0155 | 204.0 | 64.15% | 27146.5 | 55.16% | motif file (matrix) | svg |
